# Identification of a novel nidovirus in the common chameleon (*Chamaeleo chamaeleon recticrista* [Boettger, 1880])

**DOI:** 10.1101/2025.10.14.682209

**Authors:** Hana Sasaki, Sae Kawano, Mai Kishimoto, Masayuki Horie

## Abstract

Nidoviruses (the order *Nidovirales*) are positive strand RNA viruses, and the order currently contains 14 families, 48 genera, and 130 species. Novel nidoviruses have been identified in a variety of animal hosts, but many phylogenetic gaps remain, suggesting the presence of yet-to-be-identified viruses. In this study, we identified a novel nidovirus from publicly available RNA-seq data obtained from the blood of the common chameleon (*Chamaeleo chamaeleon recticrista* [Boettger, 1880]), and tentatively named it common chameleon nidovirus 1 (CCNV-1). Phylogenetic analysis showed that CCNV-1 is clustered with a member of the family *Nanhypoviridae* and some unclassified viruses designated as arteriviruses. Further, CCNV-1 genetically distinct from known viruses, exhibiting 50.9% amino acid identity in the RdRp sequence to its closest relative, Jingmen rodent arterivirus 1. Thus, this study provides novel insights into the diversity of nidoviruses.

## Introduction

Nidoviruses (the order *Nidovirales*) are positive strand RNA viruses, and the order currently contains 14 families, 48 genera, and 130 species that have been apooroved by ICTV^1^. Recent advances in high-throughput sequencing technology have identified numerous novel nidoviruses, which infect not only mammals but also reptiles^2^, amphibians^3^, fish^4^, arthropods^5^, and flatworms^6^. However, many phylogenetic gaps remain among known nidoviruses, which suggests the existence of undiscovered viruses filling the gaps. In particular, although the number of nidovirus detections in reptiles has been increasing in recent years, most studies have focused on captive individuals^2^. This suggests that a substantial number of nidoviruses remain undiscovered in wild reptile populations. Therefore, further exploration of nidoviruses is crucial for a more comprehensive understanding of their evolutionary history.

Some nidoviruses cause severe diseases in reptiles. For example, Ball python nidovirus has caused respiratory disease in captive ball pythons and is often fatal^7^. Furthermore, in 2015, a mass mortality event in the endangered wild Bellinger River snapping turtle *(Myuchelys georgesi* [Cann, 1997]) was most likely caused by infection with a nidovirus, Bellinger River virus^8^. These findings suggest that nidoviruses have the potential to cause serious diseases in both captive and wild reptiles. As mentioned above, our current understanding of the diversity of nidoviruses remains limited, and the identification of novel nidoviruses is important for the control of infectious diseases.

In this study, we detected a novel nidovirus from publicly available RNA-seq data obtained from the blood of the common chameleon (*Chamaeleo chamaeleon recticrista* [Boettger, 1880]), tentatively named it common chameleon nidovirus 1 (CCNV-1), and characterized its genetic features.

## Materials and methods

### Detection of nidovirus-like contigs

RNA-seq data (accession number SRR2962874) were downloaded from the NCBI SRA database^9^. The reads were preprocessed by fastp v0.23.2^10^ using the “-x -y -l 35” options, which were used for *de novo* assembly by rnaviralSPAdes v3.15.5^11^ and Trinity v2.15.2^12^, both with default settings. Contigs that were 5000 nucleotides or more were extracted using SeqKit v2.3.0^13^ which were used for a two-step sequence similarity search as follows.

First, a sequence similarity search was conducted against a custom database containing protein sequences of viruses belonging to the order *Nidovirales* by MMseqs2 version c48da9d781b81804727b5cccfed7f97cfcc20c9d^14^ using the assembled contigs as queries. The custom database was made as follows: amino acid sequences of viruses of the order *Nidovirales* (NCBI taxid: 76804) were downloaded from NCBI, and sequences of more than 100 amino acids were extracted by SeqKit. Then the extracted sequences were clustered by CD-HIT version 4.8.1^15^ with a threshold of 0.90. After the MMseqs2 search, the 25 contigs that had hit(s) were extracted.

The second sequence similarity search was conducted against the NCBI nr database by BLASTx (BLAST+ 2.15.0^16^) with the options “-evalue 1e-4 -word_size 3” using the extracted contigs as queries. Sequences whose BLAST best hits were viruses were extracted and manually analyzed.

To validate the accuracy of CCNV-1 contigs, the original short reads (SRR2962874) were mapped back to the CCNV-1 contigs by HISAT2 v2.2.1^17^, and the read depth at each position was calculated using SAMtools v1.16.1^18^. We defined sequences covered by at least five reads as reliable positions.

The CCNV-1 sequence was deposited to DDBJ under the accession number BR002551.

### Annotation of CCNV-1 and closely related viruses

Using Geneious Prime v2024.0 (https://www.geneious.com/), the “Find ORFs” function was applied to identify ORF sequences of CCNV-1. The slippery sequences between ORF1a and ORF1b were determined with reference to the nidovirus consensus sequence 5’-UUUAAAC-3’^19,20^. For each ORF, a domain search of amino acid sequence was performed using CD-search^21^ with the CDD v3.21–62456 PSSM database, and the E-value threshold was set to 0.01. Based on the results, annotations were added for the full-length RNA-dependent RNA polymerase (RdRp, accession cl40470), Nidoviral uridylate-specific endoribonuclease (NendoU, accession cl41717 and cl41718), 2’-O-MTase (accession cl41719), Zinc-binding domain of helicase (ZBD, accession cl41714), and Superfamily 1 RNA helicase (HEL1, accession cl28899, cl41748, and cl26263).

### Phylogenetic analyses

To collect known viral sequences for phylogenetic analysis, BLASTp search was performed against the protein sequences of viruses (taxid 10239) in the NCBI nr database on the BLAST web server, using the amino acid sequence of CCNV-1 RdRp as the query. Among the top 100 BLASTp hits, sequences with a query cover of 70% or more were extracted. The extracted sequences were clustered using CD-HIT v4.8.1 with a similarity threshold of 0.90. Finally, 73 sequences were obtained (Supplementary Table 1), which were used in the following phylogenetic analysis.

Sequences of 73 viruses closely related to CCNV-1 and 8 representative arteriviruses (Supplementary Table 1) were downloaded from NCBI. The 82 downloaded sequences and CCNV-1 were aligned using MAFFT v7.490^22^ using the L-INS-i algorithm. Based on the annotations assigned to CCNV-1, the region corresponding to the RdRp sequence was extracted from the alignment and used for phylogenetic analysis.

Phylogenetic analysis of the RdRp alignment was performed using the maximum likelihood method in MEGA v10.2.6^23^. The analysis was performed with the LG+G+I model (selected based on AICc after model testing in MEGA) and the “partial deletion (95%)” option. The reliability of the tree was assessed by 100 bootstrap resampling.

The multiple sequence alignment used in the phylogenetic analysis is available in Supplementary Material (Supplementary File 1).

### Screening reptile-derived RNA-seq data to detect CCNV-1

To identify reptile-derived RNA-seq data containing CCNV-1-derived reads, 6,549 RNA-seq datasets of Reptilia (Table S2) were mapped to the CCNV-1 sequence using Magic-BLAST 1.6.0^24^ with the options “-word_size 16 -splice F -no_unaligned.” Mapped read numbers were counted by SAMtools, and the mapping results were manually inspected using Integrative Genomics Viewer (IGV)^25^ to exclude false-positives caused by non-specific mapping..

To obtain CCNV-1-like contigs, the CCNV-1-positive RNA-seq datasets (SRR988128 and SRR768321) were downloaded. The reads were preprocessed with fastp, followed by *de novo* assembly with rnaviralSPAdes and Trinity. Then, MMseqs2 search was performed against a custom database containing protein sequences of CCNV-1 and protein sequences and other nidoviruses using the obtained contigs as queries. Contigs that had a MMseqs2 hit(s) were extracted, which were used as queries for BLASTx search against the NCBI nr database. The same options as in the identification of CCNV-1 from SRR2962874 were applied to this sequence similarity search. The obtained CCNV-1-like contigs were verified by back mapping.

## Results

### Identification of a novel nidovirus

In our previous study, we conducted large-scale viral metagenomic analyses and detected RNA viruses, including lyssaviruses and filoviruses, from publicly available reptile RNA-seq data obtained from reptiles^26,27^. However, viruses other than the above ones have not been analyzed in detail. In this study, we reanalyzed the data obtained in our previous research and focused on nidovirus-like sequences from RNA-seq data (SRR2962874) derived from the blood of the common chameleon (*Chamaeleo chamaeleon recticrista* [Boettger, 1880]).

To reanalyze the previously obtained data, *de novo* assembly and a two-step sequence similarity search were performed on the RNA-seq data (SRR2962874) described above. As a result, a nidovirus-like contig was detected, with Jingmen rodent arterivirus 1 (JRAV-1, accession number OQ715735.1, 50.9% amino acid identity in the RdRp region) as its top BLASTx hit against the NCBI nr database.

To validate the accuracy of the obtained nidovirus-like contigs, we mapped back the original sequencing reads to the contigs. In this study, we defined positions covered by at least five reads as reliable positions. As a result, a contig of 21,788 nucleotides was obtained (Fig. 1). The mapping pattern shows higher coverage depth at the 3’ end than at the 5’ end, which may reflect expression of subgenomic RNA, a hallmark of nidoviruses.

**Fig. 1.**
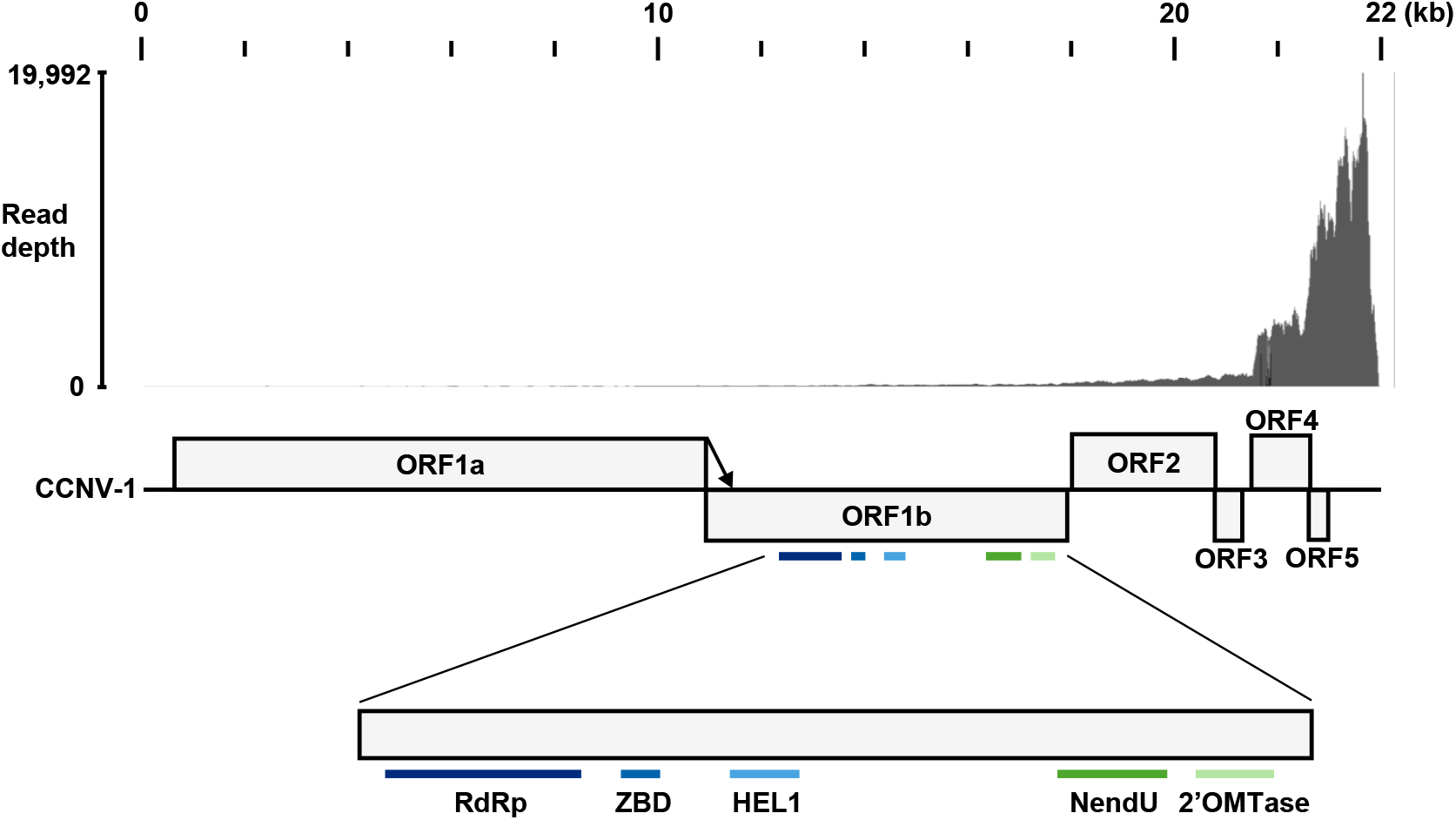
Genomic organization of common chameleon nidovirus 1. Mapped read depth (upper panel) and genome organization (lower pannel) of common chameleon nidovirus 1 (CCNV1-1) is shown. (Upper) Short reads from SRR2962874 were mapped to the CCNV-1 contigs. (Lower) Boxes indicate ORFs. Colored bars below magnified ORF1b represent domains detected by NCBI conserved domain search: dark blue, RNA-dependent RNA polymerase domain (RdRp); blue, zinc-binding domain (ZBD); light blue, helicase domain (HEL1); green, Nidoviral uridylate-specific endoribonuclease domain (NendoU); light green, 2’-O-methyltransferase (2’-O-MTase). Arrow indicates the ribosomal frameshift.

### Characterization of the nidovirus-like contigs

To determine the genomic structure of CCNV-1, we searched for ORFs on the viral contig, showing the presence of ORF1a, ORF1b, ORF2, ORF3, ORF4, and ORF5, similar to those found in other nidoviruses. A consensus slippery sequence of nidoviruses, 5’-UUUAAAC-3’, was located immediately upstream of the stop codon of ORF1a, which allows the ribosome to shift reading frames^19,20^. Assuming that the frameshift occurs in CCNV-1, translation of ORF1a may not terminate but instead continue into ORF1b, resulting in the production of a fusion protein (Fig. 1).

We next performed a functional domain search on ORF1a, ORF1b, ORF2, ORF3, ORF4, and ORF5. As a result of the domain search using CD-search, five domains were detected within ORF1b: RdRp, NendoU, 2’-O-MTase, ZBD, and HEL1 (Fig. 1 and Supplementary Table 2). Among these, the first four domains were identified in full length, whereas the HEL1 domain was only partially identified. No functional domains were detected within ORF1a, ORF2, ORF3, ORF4, or ORF5.

### Phylogenetic analysis

To infer the evolutionary relationships between CCNV-1 and other nidoviruses, we performed phylogenetic analyses using the deduced amino acid sequences of the RdRp domain. The analysis revealed that CCNV-1 clustered with Wuhan Japanese halfbeak arterivirus (WJHAV, accession number MG600020.1), a member of *Nanhypoviridae* (Fig. 2). Note that this cluster also included three additional viruses classified as arteriviruses in NCBI GenBank: Hainan hebius popei arterivirus (HHPAV, accession number MG600021.1), JRAV-1, and *Arteriviridae* sp. (accession number MT085142.1) (see Discussion).

**Fig. 2.**
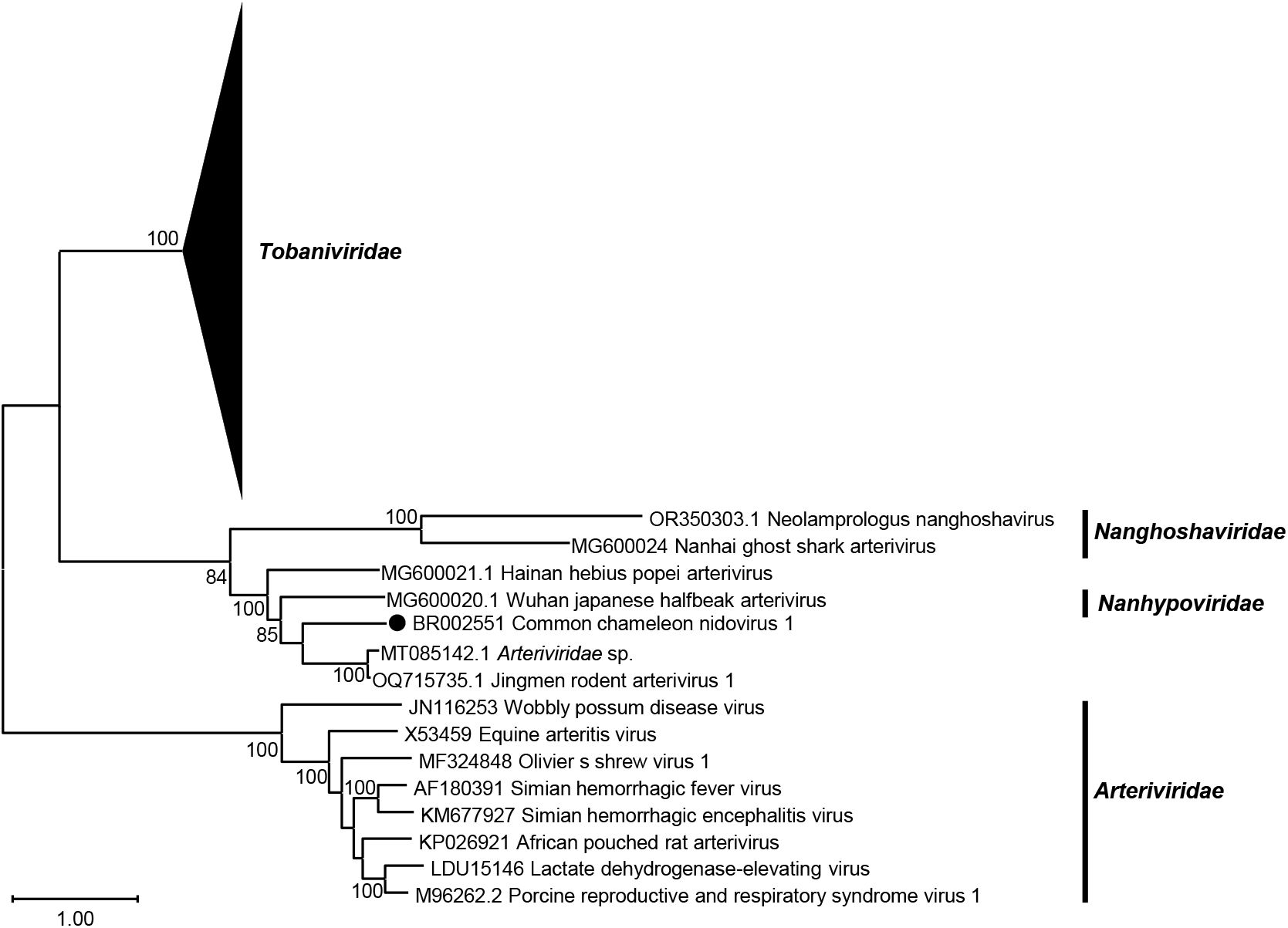
Phylogenetic relationship between common chameleon nidovirus 1 and nidoviruses. Phylogenetic relationship among viruses from the families of *Nanghoshaviridae, Nanhypoviridae* and *Arteriviridae, Tobaniviridae*. Phylogenetic tree was inferred by the maximum likelihood method using amino acid sequences of the RdRp domein. The scale bar indicates the number of amino acid substitutions per site. Bootstrap values less than 80 were omitted from this tree.

### Characterization of closely related viruses

To compare the genome structures of viruses belonging to the same cluster with CCNV-1, we analyzed the ORFs on their genomes (Fig. 3). In HHPAV and WJHAV, ORF1ab, ORF2, ORF3, and ORF4 were identified. In JRAV-1, ORF1ab, ORF2, ORF3, ORF4, and ORF5 were identified. For *Arteriviridae* sp., only a partial sequence corresponding to a portion of ORF1b was identified. Furthermore, the slippery sequence 5’-UUUAAAC-3’ was detected immediately upstream of ORF1a in HHPAV, JRAV-1, and WJHAV, similar to CCNV-1.

**Fig. 3.**
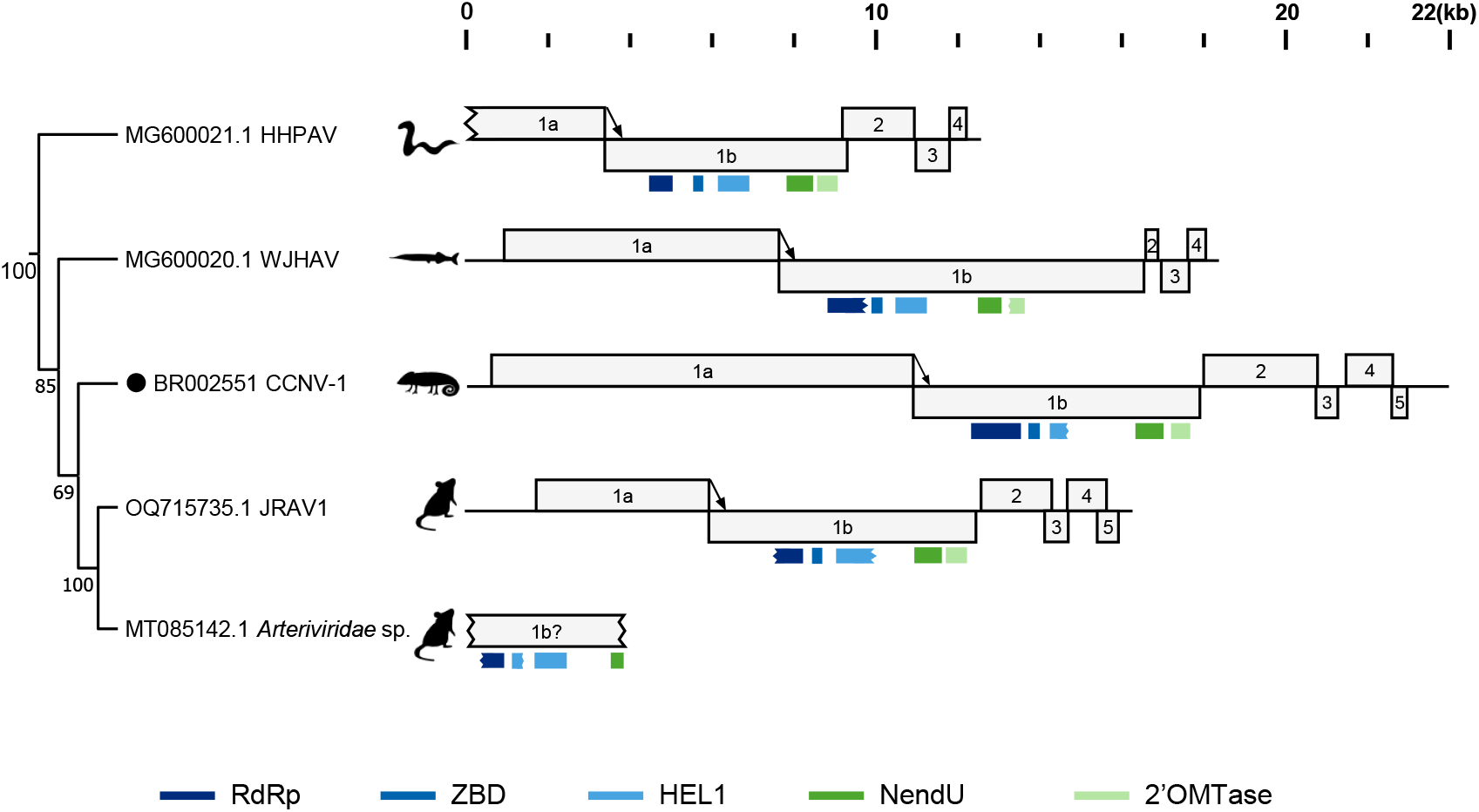
Comparison of genome structure of common chameleon nidovirus 1 and related viruses. Phylogenetic tree (only topology) and genome organizations of the shown viruses. The topology of the three is adopted from Fig. 1. Colored bars below magnified ORF1b represent domains detected by NCBI conserved domain search: dark blue, RNA-dependent RNA polymerase domain (RdRp); blue, zinc-binding domain (ZBD); light blue, helicase domain (HEL1); green, Nidoviral uridylate-specific endoribonuclease domain (NendoU); light green, 2’-O-methyltransferase (2’-O-MTase). Partially identified domains are shown as jagged lines.

We also performed a functional domain search within the ORFs of those viruses. We identified the following conserved domains within ORF1b: RdRp, NendoU, 2’-O-MTase, HEL1, and ZBD (Fig. 3 and Supplementary Table 2). In HHPAV, all five domains were detected as nearly complete. In WJHAV, ZBD and HEL1 were detected as nearly complete, whereas RdRp, NendoU, and 2’-O-MTase were only partially detected. In JRAV-1, ZBD, HEL1, NendoU, and 2’-O-MTase were detected as nearly complete, while RdRp was only partially detected. In *Arteriviridae* sp., HEL1, NendoU were detected as nearly complete, whereas RdRp and ZBD were only partially detected. We also analyzed ORFs other than ORF1b in each virus, but no functional domains were identified.

### CCNV-1 was exclusively detected from the common chameleon

To further detect CCNV-1, we mapped 6,549 reptile RNA-seq datasets to CCNV-1. We found two additional datasets (SRR988128 and SRR768321) contain reads mappable to CCNV-1 (Fig. 4 and Table 1), which originate from the blood and multi-tissue samples (the brain, lungs, skeletal muscle, and heart) of *C. chamaeleon*, respectively. From the dataset SRR988128, we obtained a contig of nearly complete genome of CCNV-1 (20,243 nt), showing 96.4% nucleotide identity to the CCNV-1 sequence detected in SRR2962874.

**Table 1.**
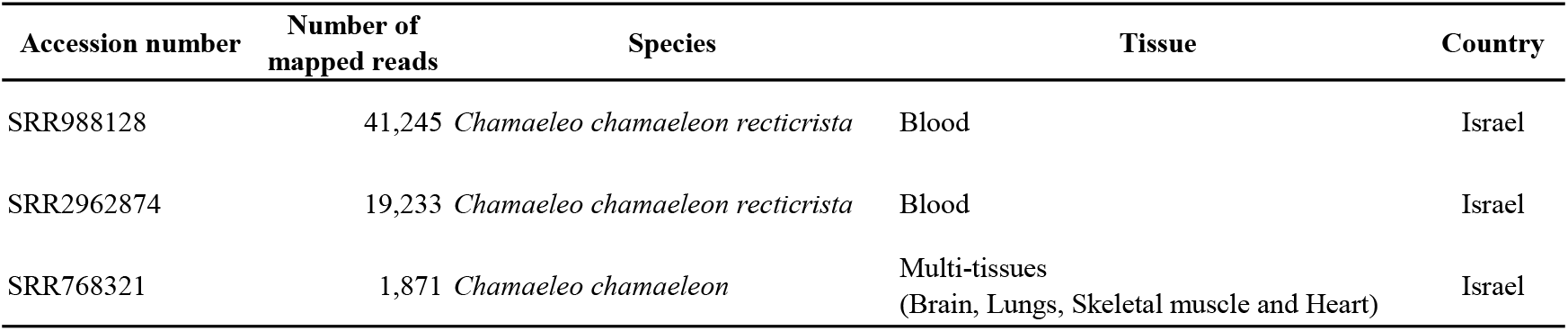
List of RNA-seq datasets containing common chameleon nidovirus 1-derived reads.

| Accession number | Number of mapped reads | Species | Tissue | Country |
| --- | --- | --- | --- | --- |
| SRR988128 | 41,245 | *Chamaeleo chamaeleon recticrista* | Blood | Israel |
| SRR2962874 | 19,233 | *Chamaeleo chamaeleon recticrista* | Blood | Israel |
| SRR768321 | 1,871 | *Chamaeleo chamaeleon* | Multi-tissues<br>(Brain, Lungs, Skeletal muscle and Heart) | Israel |

**Fig. 4.**
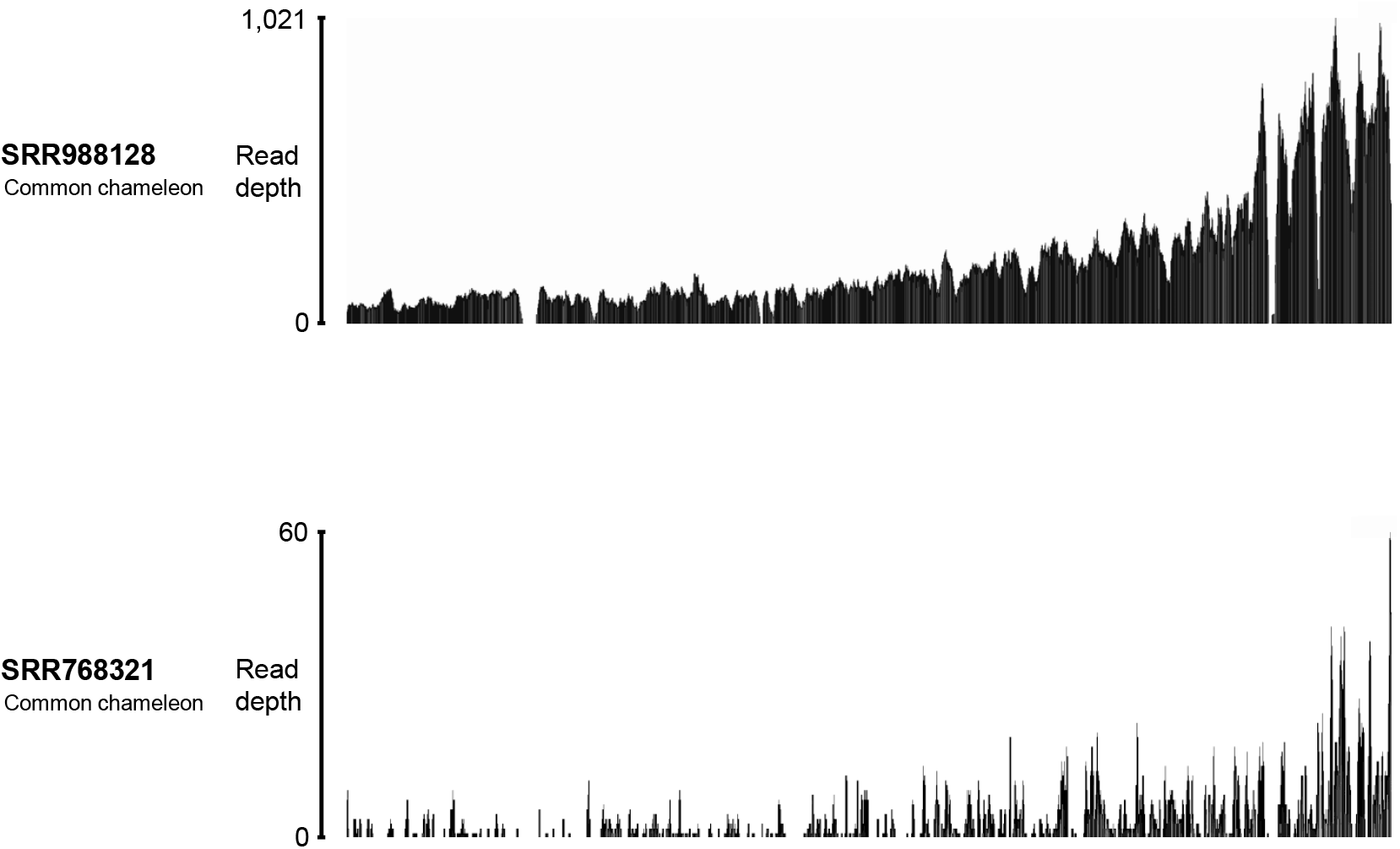
Mapping of RNA-seq datasets (SRR988128 and SRR768321) to CCNV-1. Short reads from SRR988128 and SRR768321 were mapped to CCNV-1 and mapped read depths of are visualized.

## Discussion

A wide variety of nidoviruses from diverse hosts have been identified thus far. However, many phylogenetic gaps are still present, suggesting the existence of undiscovered nidoviruses that may fill these gaps. In this study, we identified a novel nidovirus from publicly available RNA-seq data (SRR2962874) obtained from the blood of the common chameleon, tentatively named CCNV-1. The phylogenetic analysis showed CCNV-1 clustered with viruses belonging to *Nanhypoviridae* and classified as arteriviruses in NCBI. However, it was genetically distinct from the other viruses in the cluster, suggesting that it may represent a novel virus species (see below for details). Thus, this study contributes to a deeper understanding of the diversity and evolution of nidoviruses.

This study strongly suggests that CCNV-1, HHPAV, JRAV-1, and Arteriviridae sp. (MT085142.1) are members of the family *Nanhypoviridae* and that the names of HHPAV, WJHAV, JRAV-1, and Arteriviridae sp. may require renaming. CCNV-1, HHPAV, JRAV-1, and Arteriviridae sp. (MT085142.1) clustered with WJHAV, an ICTV approved member of the family *Nanhypoviridae* (Fig. 2). Although HHPAV, WJHAV, JRAV-1, and Arteriviridae sp. are currently named as “arteriviruses,” they are phylogenetically distinct from known members of the family *Arteriviridae* (Fig. 2). In addition, domain searches for CCNV-1, HHPAV, WJHAV, and JRAV-1 detected 2’-O-MTase. Although this domain is known to be absent in viruses of the family *Arterivirida* ^28^, all four viruses contain it, strongly suggesting that they are structurally distinct from the members of the family *Arteriviridae*. Assigning the same name to viruses with significant differences in molecular structure and phylogenetic relationships may compromise the accuracy of future research. Therefore, viral naming should be revised appropriately, based on evolutionary background and characteristics.

In this study, we performed phylogenetic analysis (Fig. 2) using only the amino acid sequence of the RdRp domain, which may compromise the robustness of the tree. In this study, we tried to identify domains conserved among nidoviruses on the ORF1ab proteins of CCNV-1 and the other related viruses, but we could not identify complete length of those domains. Therefore, we used only RdRp to infer phylogenetic relationship among those viruses. However, some internal branches were not well supported. The RdRp domain is found in all nidoviruses and is useful for phylogenetic analysis, but its short length could limit detailed classification. To infer evolutionary relationship more robustly, it may be helpful to identify and include other conserved domains. 3CLpro, NiRAN, RdRp, ZBD, and HEL1 are listed by the ICTV as criteria for species demarcation within the family *Arteriviridae*^29^. Therefore, by reanalyzing with ZBD, HEL1, the 3CLpro, and NiRAN in addition to the RdRp used in this study, the phylogenetic relationship between CCNV-1 and the related viruses may be inferred more accurately.

Our study may suggest host specificity of CCNV-1 for the common chameleon. Across 6,549 reptile RNA-seq datasets, CCNV-1-derived reads were exclusively detected in the common chameleon, suggesting the host specificity of CCNV-1. However, due to the limited sample sizes and taxonomic biases of publicly available datasets, we cannot exclude the possibility that CCNV-1 infection in other reptiles could have been missed. Future studies with broader taxonomic sampling and analysis will clarify the host range of CCNV-1 more accurately.

Taken together, this study filled a phylogenetic gap within nidoviruses, thereby expanding our understanding of their diversity and evolution. However, the phylogenetic gaps among nidoviruses remain large. Furthermore, their modes of transmission, pathogenicity, replication and proliferation mechanisms, and evolutionary history of nidoviruses remain largely unclear. Further studies, including the identification of new hosts and the accumulation of additional sequence data, are needed to better understand the diversity of nidoviruses.

## Acknowledgements

The super-computing resources were provided by Human Genome Center, the Institute of Medical Science, the University of Tokyo. This study was supported by KAKENHI grant numbers 19H04833 (MH), 21H01199 (MH), 23K20902 (MH), and 24K21922 (MH) and Osaka Metropolitan University (OMU) Strategic Research Promotion Project (Young Researcher; MK).

## References

1. Simmonds, P. et al. Changes to virus taxonomy and the ICTV Statutes ratified by the International Committee on Taxonomy of Viruses (2024). Arch. Virol. 169, 236 (2024).

2. Parrish, K., Kirkland, P. D., Skerratt, L. F. & Ariel, E. Nidoviruses in Reptiles: A Review. Front. Vet. Sci. 8, 733404 (2021).

3. Lauber, C. et al. Deep mining of the Sequence Read Archive reveals major genetic innovations in coronaviruses and other nidoviruses of aquatic vertebrates. PLoS Pathog. 20, e1012163 (2024).

4. Schütze, H. et al. Characterization of White bream virus reveals a novel genetic cluster of nidoviruses. J. Virol. 80, 11598–11609 (2006).

5. Nga, P. T. et al. Discovery of the first insect nidovirus, a missing evolutionary link in the emergence of the largest RNA virus genomes. PLoS Pathog. 7, e1002215 (2011).

6. Saberi, A., Gulyaeva, A. A., Brubacher, J. L., Newmark, P. A. & Gorbalenya, A. E. A planarian nidovirus expands the limits of RNA genome size. PLoS Pathog. 14, e1007314 (2018).

7. Hoon-Hanks, L. L. et al. Respiratory disease in ball pythons (Python regius) experimentally infected with ball python nidovirus. Virology 517, 77–87 (2018).

8. Zhang J. et al. Identification of a novel nidovirus as a potential cause of large scale mortalities in the endangered Bellinger River snapping turtle (Myuchelys georgesi). PLOS ONE 13, e0205209 (2018).

9. Sayers, E. W. et al. Database resources of the National Center for Biotechnology Information. Nucleic Acids Res. 52, D33–D43 (2024).

10. Chen, S., Zhou, Y., Chen, Y. & Gu, J. fastp: an ultra-fast all-in-one FASTQ preprocessor. Bioinforma. Oxf. Engl. 34, i884–i890 (2018).

11. Bushmanova, E., Antipov, D., Lapidus, A. & Prjibelski, A. D. rnaSPAdes: a de novo transcriptome assembler and its application to RNA-Seq data. GigaScience 8, giz100 (2019).

12. Grabherr, M. G. et al. Full-length transcriptome assembly from RNA-Seq data without a reference genome. Nat. Biotechnol. 29, 644–652 (2011).

13. Shen, W., Le, S., Li, Y. & Hu, F. SeqKit: A Cross-Platform and Ultrafast Toolkit for FASTA/Q File Manipulation. PloS One 11, e0163962 (2016).

14. Steinegger, M. & Söding, J. MMseqs2 enables sensitive protein sequence searching for the analysis of massive data sets. Nat. Biotechnol. 35, 1026–1028 (2017).

15. Fu, L., Niu, B., Zhu, Z., Wu, S. & Li, W. CD-HIT: accelerated for clustering the next-generation sequencing data. Bioinforma. Oxf. Engl. 28, 3150–3152 (2012).

16. Camacho, C. et al. BLAST+: architecture and applications. BMC Bioinformatics 10, 421 (2009).

17. Kim, D., Paggi, J. M., Park, C., Bennett, C. & Salzberg, S. L. Graph-based genome alignment and genotyping with HISAT2 and HISAT-genotype. Nat. Biotechnol. 37, 907–915 (2019).

18. Danecek, P. et al. Twelve years of SAMtools and BCFtools. GigaScience 10, giab008 (2021).

19. Schütze, H. et al. Characterization of White Bream Virus Reveals a Novel Genetic Cluster of Nidoviruses. J. Virol. 80, 11598–11609 (2006).

20. Brierley, I., Digard, P. & Inglis, S. C. Characterization of an efficient coronavirus ribosomal frameshifting signal: Requirement for an RNA pseudoknot. Cell 57, 537–547 (1989).

21. Marchler-Bauer, A. & Bryant, S. H. CD-Search: protein domain annotations on the fly. Nucleic Acids Res. 32, W327–331 (2004).

22. Katoh, K. & Standley, D. M. MAFFT multiple sequence alignment software version 7: improvements in performance and usability. Mol. Biol. Evol. 30, 772–780 (2013).

23. Kumar, S., Stecher, G., Li, M., Knyaz, C. & Tamura, K. MEGA X: Molecular Evolutionary Genetics Analysis across Computing Platforms. Mol. Biol. Evol. 35, 1547–1549 (2018).

24. Boratyn, G. M., Thierry-Mieg, J., Thierry-Mieg, D., Busby, B. & Madden, T. L. Magic-BLAST, an accurate RNA-seq aligner for long and short reads. BMC Bioinformatics 20, 405 (2019).

25. Robinson, J. T. et al. Integrative genomics viewer. Nat. Biotechnol. 29, 24–26 (2011).

26. Horie, M., Akashi, H., Kawata, M. & Tomonaga, K. Identification of a reptile lyssavirus in Anolis allogus provided novel insights into lyssavirus evolution. Virus Genes 57, 40–49 (2021).

27. Horie, M. Identification of a novel filovirus in a common lancehead (Bothrops atrox (Linnaeus, 1758)). J. Vet. Med. Sci. 83, 1485–1488 (2021).

28. Lehmann, K. C. et al. Arterivirus nsp12 versus the coronavirus nsp16 2’-O-methyltransferase: comparison of the C-terminal cleavage products of two nidovirus pp1ab polyproteins. J. Gen. Virol. 96, 2643–2655 (2015).

29. Brinton, M. A. et al. ICTV Virus Taxonomy Profile: Arteriviridae 2021. J. Gen. Virol. 102, 001632 (2021).

